# PPARG directs trophoblast cell fate and establishment of the uterine-placental interface

**DOI:** 10.1101/2025.10.08.681177

**Authors:** Esteban M. Dominguez, Ayelen Moreno-Irusta, Khursheed Iqbal, Keting Chen, Alex Finlinson, Marc Parrish, Hiroaki Okae, Takahiro Arima, Geetu Tuteja, Michael J. Soares

## Abstract

The expansion and differentiation of trophoblast stem (**TS**) cells are critical for defining fundamental properties of the placenta. Specialized trophoblast cells exit the placenta and enter and transform the uterus, including restructuring uterine spiral arteries. In the human these cells are named extravillous trophoblast (**EVT**) cells, whereas in the rat they are termed invasive trophoblast cells. Mechanisms governing invasive trophoblast cell differentiation remain poorly understood. We investigated peroxisome proliferator-activated receptor gamma (**PPARG**) as a potential regulator of EVT/invasive trophoblast cell development. In first trimester human placentas, PPARG was expressed in the EVT cell column and increased in amount as human TS cells differentiated into EVT cells. PPARG disruption impaired EVT cell differentiation. Rat invasive trophoblast cells similarly expressed PPARG. Conditional inactivation of PPARG within rat invasive trophoblast cells was used to assess the in vivo role of PPARG on the uterine-placental interface. PPARG was established as an essential cell-autonomous regulator of the invasive trophoblast cells. In conclusion, PPARG is a conserved regulator of placentation and is essential for directing trophoblast cell-guided uterine transformation.

## Introduction

The uterine-placental interface is a dynamic site where uterine and placental structures cooperate to establish a protective environment that redirects maternal resources to support embryonic and fetal development (Georgiades *et al*, 2002; Maltepe & Fisher, 2015). Both humans and rats possess a hemochorial placenta, characterized by deep trophoblast cell invasion into the uterine parenchyma and remodeling of the uterine vasculature to ensure optimal maternal blood supply to the placenta (Pijnenborg *et al*, 1981; Soares *et al*, 2012). This vascular transformation is critical for successful pregnancy (Pijnenborg *et al*, 2006; Aye *et al*, 2025). Impairments in this process can lead to complications such as pregnancy loss, preeclampsia, intrauterine growth restriction, and preterm birth (Aye *et al*, 2025; Brosens *et al*, 2011; Fisher, 2015; Brosens *et al*, 2019; Harris *et al*, 2019). These complications significantly contribute to maternal and fetal morbidity and mortality (Fisher, 2015; Brosens *et al*, 2019; Kaufmann *et al*, 2003). Furthermore, suboptimal intrauterine conditions can trigger fetal adaptations that increase susceptibility to disease later in life (Calkins & Devaskar, 2011; Gluckman *et al*, 2008).

Throughout gestation, maternal blood flows into the placenta, delivering essential nutrients and oxygen to support fetal development. As pregnancy progresses, maternal uterine spiral arteries are remodeled into distended, low-resistance vessels capable of meeting the increasing demands of the growing fetus (Kaufmann *et al*, 2003; Harris, 2010). At the core of spiral artery remodeling are invasive trophoblast cells, referred to as extravillous trophoblast (**EVT**) cells in the human (Velicky *et al*, 2015). These cells take two routes into the uterus. They migrate inside the uterine vasculature and within the uterine stroma situated between uterine blood vessels and are referred to as endovascular invasive trophoblast/EVT cells and interstitial invasive trophoblast/EVT cells, respectively. Endovascular invasive trophoblast/EVT cells replace endothelial cells and adopt a pseudo-endothelial phenotype (Hemberger *et al*, 2003; Rai & Cross, 2014). Given the critical role of invasive trophoblast/EVT cells in uterine spiral artery remodeling, it is crucial to identify mechanisms that regulate invasive trophoblast/EVT cell development.

Peroxisome proliferator-activated receptor gamma (**PPARG**) is a member of the nuclear hormone receptor superfamily of ligand-activated transcription factors (Sauer, 2015). PPARG is essential for placental development in the mouse (Barak *et al*, 1999; Kubota *et al*, 1999). Deficiency of PPARG results in embryonic lethality at mid-gestation (Barak *et al*, 1999; Kubota *et al*, 1999). In addition, PPARG is expressed in the human placenta and is involved in human trophoblast cell development and regulating key placental functions (Schaiff *et al*, 2006; Tarrade, 2001; Tarrade *et al*, 2001; Pavan *et al*, 2003; Fournier *et al*, 2007; Schaiff *et al*, 2000; Shalom-Barak *et al*, 2004). Among its diverse actions on placentation is its potential involvement in regulating the invasive trophoblast cell lineage. Within rat and human placentation sites, PPARG is prominently expressed in invasive trophoblast/EVT cell lineages (Liu *et al*, 2018; Vento-Tormo *et al*, 2018; Scott *et al*, 2022). PPARG dysregulation is also evident in diseases associated with failures in trophoblast cell-guided uterine spiral artery remodeling such as preeclampsia and intrauterine growth restriction (Rodie *et al*, 2005; Waite *et al*, 2005; Holdsworth-Carson *et al*, 2010).

In this study, we investigated the involvement of PPARG on EVT cell lineage development in vitro using human trophoblast stem (**TS**) cells. Human TS cells are a robust in vitro model system and have provided new insights into the regulation of trophoblast cell differentiation (Okae *et al*, 2018; Varberg *et al*, 2023). The contributions of PPARG to invasive trophoblast cell development were also examined in vivo using a conditionally disrupted *Pparg* rat model. Our findings demonstrate that PPARG is a critical and conserved regulator of invasive trophoblast cell development and invasive trophoblast cell-guided transformation of the uterine vasculature.

## Results

### PPARG is expressed in EVT cells

Distributions of PPARG transcript and protein in first trimester human placenta tissue were determined by in situ hybridization and immunohistochemistry, respectively. PPARG transcript (Fig. 1A) and protein (Fig. 1B) were ubiquitously expressed throughout trophoblast cells within villous and extravillous compartments of the human placenta, including EVT cells. The transcript for notum, palmitoleoyl protein carboxylesterase (**NOTUM**), an EVT cell-associated transcript (Shukla *et al*, 2024), co-localized with PPARG transcripts in the distal region of the EVT cell column (Fig. 1A). PPARG protein was located within nuclei of EVT cells (Fig. 1B). These observations are consistent with previous reports (Tarrade *et al*, 2001; Waite *et al*, 2000). PPARG expression was also examined in CT27 (X,X) human TS cells, in the stem state and after EVT cell differentiation (Okae *et al*, 2018). Expression of *TEAD4* and *TP63* defined the stem state of these cells, whereas *HLA-G* and *MMP2* expression characterized EVT cell differentiation (Fig. EV1A). PPARG transcript and protein expression increased from the stem state to EVT cell differentiation state (Fig. 1C). Thus, PPARG is expressed in EVT cells of the developing human placenta and in human EVT cells following their differentiation from TS cells.

**Figure 1.**
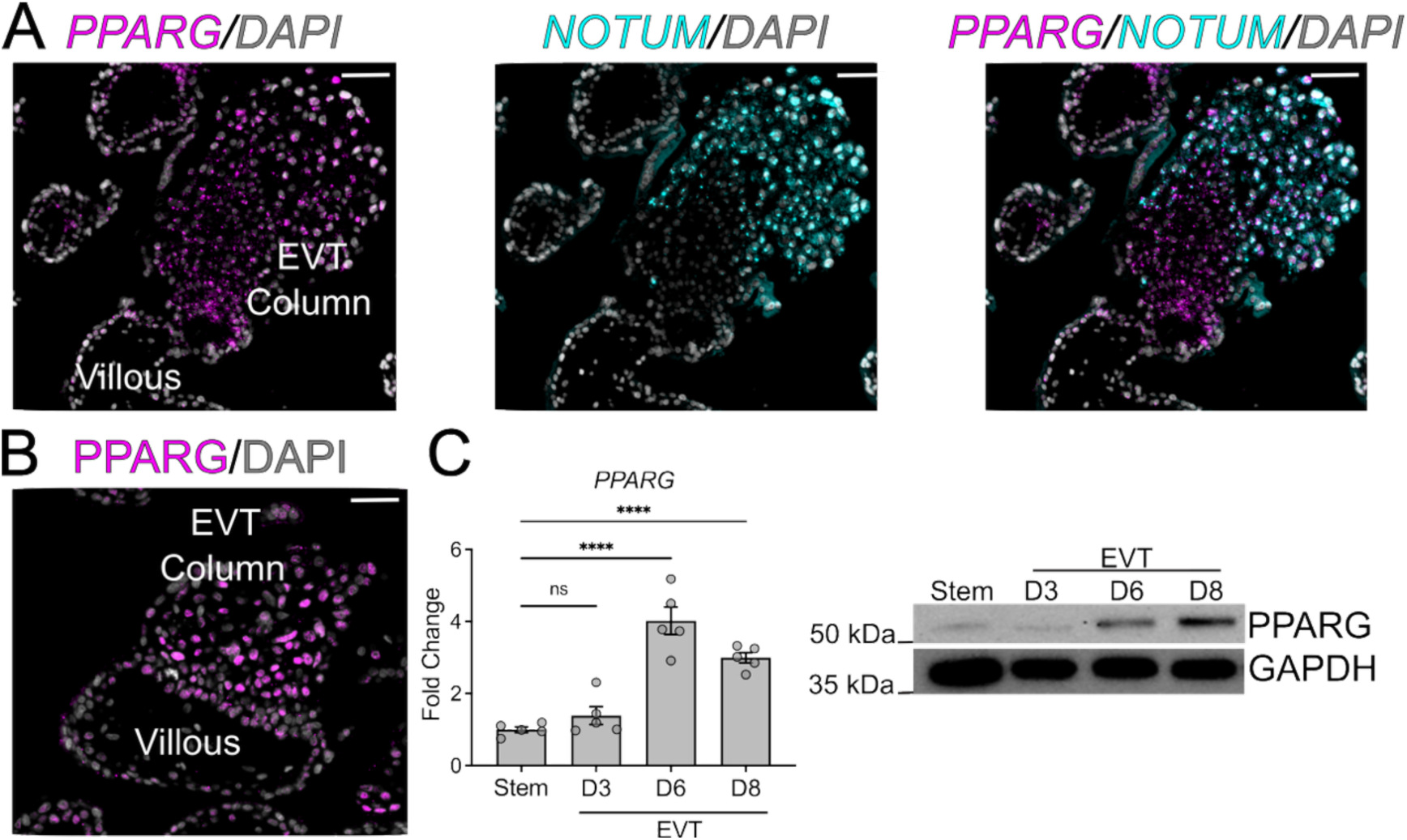
PPARG is expressed in EVT cells. **(A)** *PPARG* transcript (magenta) was localized throughout the EVT column in first trimester human placenta (12 weeks) using in situ hybridization. *NOTUM* (cyan) was used as a distal EVT cell specific transcript. **(B)** PPARG immunohistochemistry (magenta) showed nuclear localization within cells of the first trimester EVT cell column (12 weeks). Scale bars for panels A and B=500 μm. **(C)** In vitro *PPARG* transcript and protein expression in CT27 (X,X) TS cells in the stem state, day 3 (D3), day 6 (D6), and day 8 (D8) of EVT cell differentiation (n=5). Data are presented as mean ± standard error of the mean. Each data point represents a biological replicate. Statistical analysis was performed using ANOVA followed by Dunnett’s post hoc test: ****p<0.0001.

### PPARG effects on EVT cell differentiation

The expression of PPARG within EVT cells prompted an investigation into a possible role for PPARG in EVT cell development. A lentiviral-mediated short hairpin RNA (**shRNA**) loss-of-function strategy was used to inhibit PPARG in CT27 (X,X) TS cells. PPARG transcript and protein were significantly decreased in EVT cells stably expressing either of two different PPARG shRNAs when compared to expression of a control shRNA (Fig. 2A). The specificity of PPARG-targeting shRNAs was assessed by monitoring the expression of other members of the PPAR family. Expression of *PPARA* and *PPARD* transcripts were not affected following treatment with PPARG shRNAs (Fig. EV1B). Treatment with PPARG shRNAs slowed TS cell proliferation without changes in stem state associated markers TEAD4 and TP63 (Fig. EV1C-E). Disruption of PPARG affected the morphology of the EVT cell differentiation state, including a decrease in cell elongation and the presence of stem state-like cell clusters (Fig. 2B). Cells expressing PPARG shRNAs showed significantly less migration than EVT cells expressing a control shRNA (Fig. 2C). Similar findings were observed in CT29 (X,Y) TS cells following their manipulation with PPARG shRNAs (Fig. EV2A-C). Additionally, exposure to a small molecule PPARG antagonist (GW9662) similarly negatively affected stem state proliferation, EVT cell differentiation, and cell migration (Fig. EV3A-J). Collectively, the findings indicate that PPARG possesses a fundamental role in regulating EVT cell development and function.

**Figure 2.**
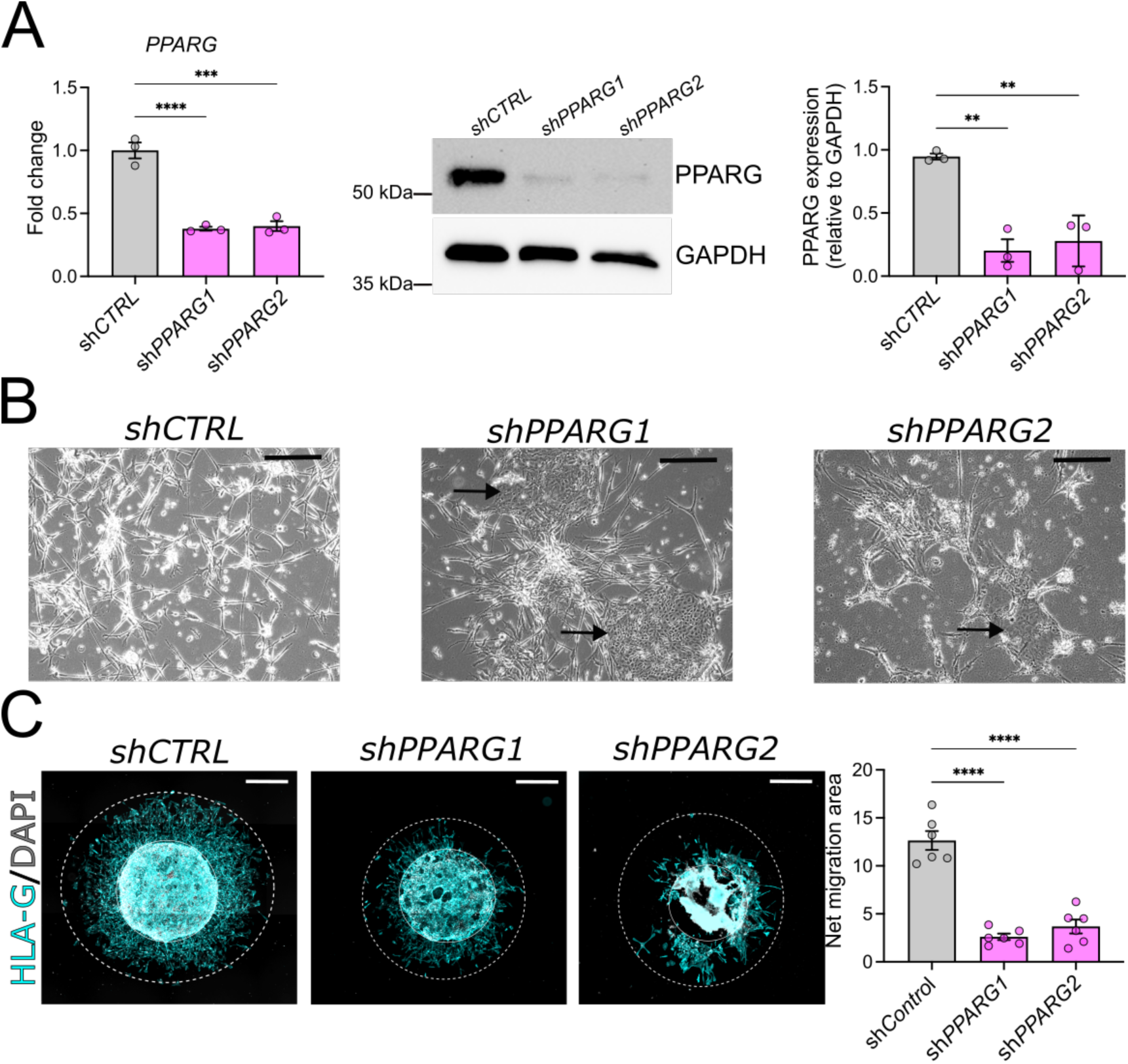
PPARG regulates EVT cell differentiation. **(A)** In vitro *PPARG* transcript and protein levels after 8 days of EVT cell differentiation in control versus PPARG1 and PPARG2 shRNAs-treated cells (n=3). **(B)** Phase-contrast images showing the morphology of EVT cells expressing control or PPARG1 and PPARG2 shRNAs. The arrow indicates a cluster of cells exhibiting a stem-like morphology, scale bar=100 μm. **(C)** HLA-G immunofluorescence detection of cell migration in control versus PPARG1 and PPARG2 shRNA-treated cells (n=6), scale bar=500 μm. Data are presented as the mean ± standard error of the mean. Each data point represents a biological replicate. Statistical analysis was performed using one-way ANOVA followed by Dunnett’s post hoc tests: ** p<0.01, ***p<0.001, ****p<0.0001.

### PPARG effects on trophoblast organoids

We next examined the involvement of PPARG in establishment and behavior of TS cell-derived trophoblast organoids. We initially validated the TS cell-derived trophoblast organoid culture system (Fig. EV 4A–D) and then examined control or PPARG deficient trophoblast organoids. The ability to form trophoblast organoids was not affected by PPARG disruption (Fig. 3A); however, PPARG inhibition did affect EVT cell differentiation from trophoblast organoids (Fig. 3B). This finding was supported by inspection of the morphology of the trophoblast organoids and their expression of TS cell stem state transcripts (*TEAD4, TFAP2C, TP63, CDH1*) and EVT cell-associated transcripts (*MMP15, HLA-G, FSTL3, ASCL2, DLX6, PLAC8*) (Fig. 3C).

**Figure 3.**
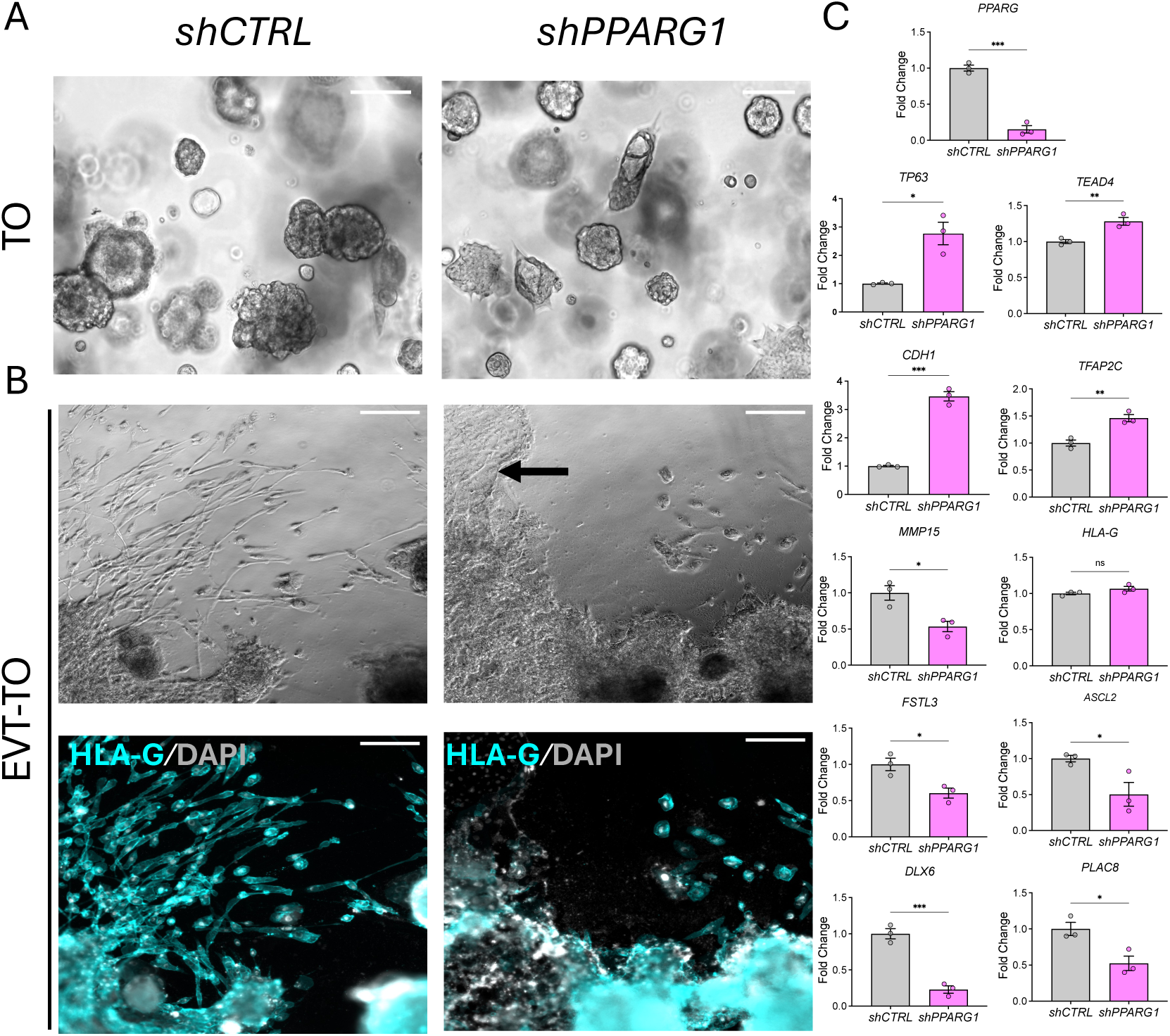
Examination of the role of PPARG in EVT cell differentiation using trophoblast organoids. **(A)** Phase-contrast images of trophoblast organoids (**TO**) expressing control (**shCTRL**) and PPARG (**shPPARG**) shRNAs, scale bar=100 μm. **(B)** Phase-contrast (top panel) and HLA-G immunostaining (bottom panel) images of shCTRL and shPPARG TO transferred to EVT cell differentiation conditions (**EVT-TO**). Arrows indicate the location of cells exhibiting stem-like morphology, scale bar=100 μm. **(C)** RT-qPCR analysis of shCTRL and shPPARG TO exposed to EVT cell differentiation conditions. Data are presented as the mean ± standard error of the mean. Each data point represents a biological replicate. Statistical analysis was performed using unpaired t-tests: *P<0.05, **p<0.01, ***p<0.001.

### Identification of PPARG gene targets in EVT cells

PPARG is a transcription factor, which prompted an examination of transcripts affected by PPARG disruption. RNA sequencing (**RNA-seq**) was performed, and transcriptomic profiles analyzed following eight days of EVT cell differentiation in control shRNA and PPARG1 shRNA treated cells (Appendix Dataset 1). A total of 5,845 differentially expressed genes were identified in control versus PPARG1 shRNA treated cells (p<0.05, and fold change of 1.5), including 2,510 upregulated and 3,335 downregulated genes. The differential gene expression profile is presented as a volcano plot (Fig. 4A). Selected upregulated and downregulated genes connected to trophoblast cell biology are shown in a heat map (Fig. 4B). We identified upregulation of several genes associated with the TS cell stem state (e.g. *TEAD4, TFAP2C, TP63, CDH1*) and downregulation of several EVT cell-associated genes (e.g. *ITGA1, ITGAV, FLT1, EPAS1, DLX6, PLAC8, TFPI, FSTL3*). These findings were independently validated by reverse transcriptase-quantitative polymerase chain reaction (**RT-qPCR,** Fig. 4C, Appendix Fig.1). Gene ontology (**GO**) (Fig. 4D and Appendix Dataset 2,3) and gene set enrichment analysis (**GSEA**) (Fig. 4E and Appendix Dataset 4) implicated PPARG in the upregulation of cell cycle associated genes and the downregulation of cell projection assembly associated genes. The latter cellular process directly connects PPARG with cell elongation and motility that are acquired as TS cells transit from the stem state to EVT cells.

**Figure 4.**
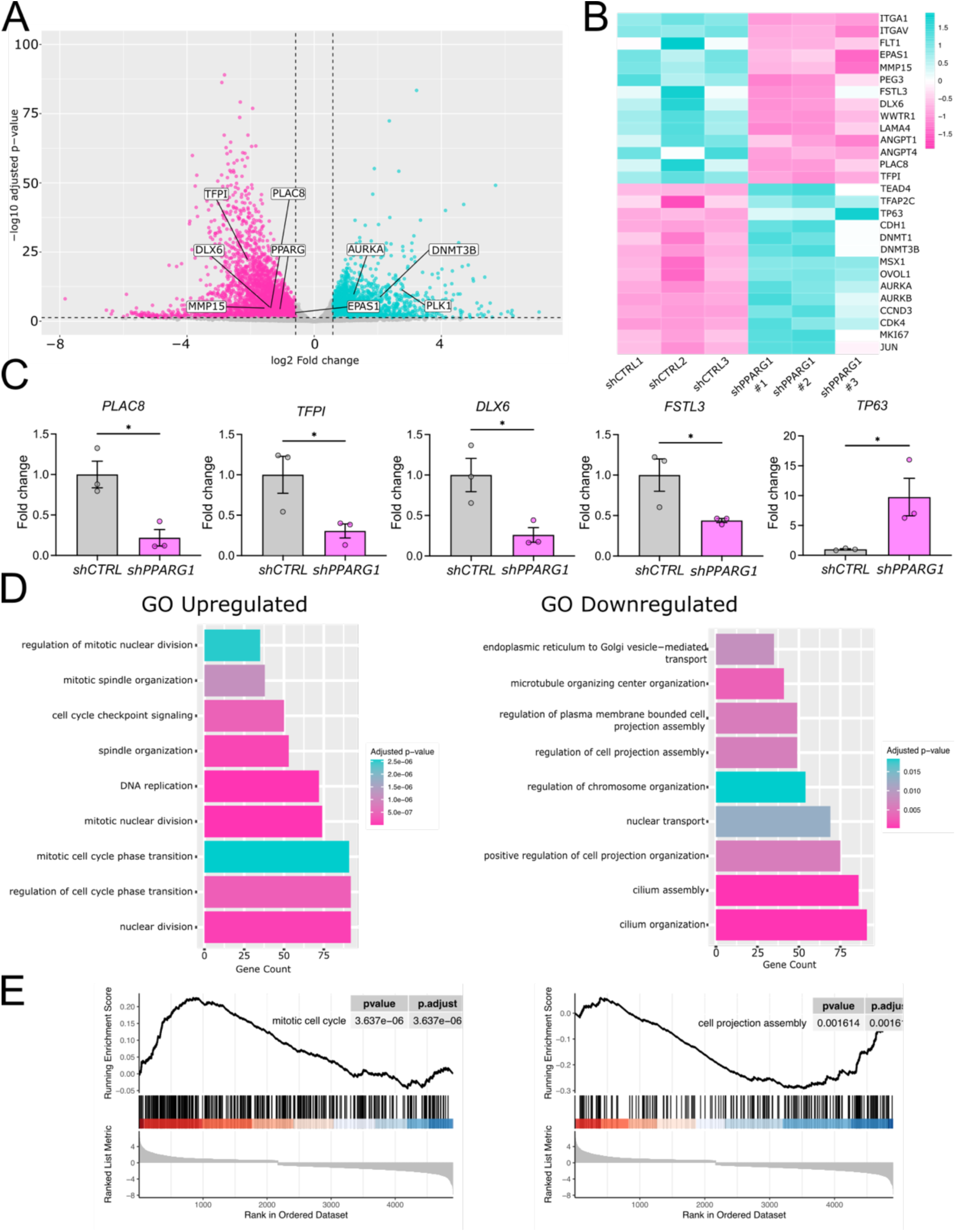
Effects of PPARG on the EVT cell transcriptome. **(A)** Volcano plot and **(B)** heat map depicting RNA-seq analysis of control shRNA and PPARG shRNA for cells exposed to culture conditions used to promote EVT cell differentiation (n=3 per group). Downregulated transcripts are represented by magenta dots (p<0.05, fold change <1.5) and cyan dots represent upregulated transcripts (p<0.05, fold change>1.5). Z-scores of totals read values are represented in the heat map. **(C)** RT-qPCR validation of putative PPARG targets in control and PPARG shRNA treated cells. **(D)** Gene ontology (GO) analyses of upregulated and downregulated genes in control and PPARG shRNA treated TS cells exposed to culture conditions used to promote EVT cell differentiation. **(E)** Gene set enrichment analyses of upregulated and downregulated genes in control and PPARG shRNA treated TS cells exposed to culture conditions used to promote EVT cell differentiation. CT27 (X,X) TS cell line was used for all experiments shown. Data are presented as the mean ± standard error of the mean. Each data point represents a biological replicate. Statistical analyses were performed using unpaired t-tests: *p<0.05.

Since, PPARG is a transcription factor, we next analyzed genome-wide PPARG-DNA interactions using chromatin immunoprecipitation sequencing (**ChIP-seq**). We identified 7,319 genes with at least one PPARG bound region (peak) in differentiated EVT cells (Appendix Dataset 5). The two most prevalent motifs identified within the peaks were known PPARG DNA binding motifs (peroxisome proliferator response element, **PPRE**) and primarily consisted of a direct repeat 1 (**DR1**) sequence (Fig. 5A). Most PPARG-DNA peaks (∼7000) were located within introns or intergenic regions of protein-coding genes (Fig. 5B,C), suggesting the involvement of PPARG in the regulation of enhancer activity. Integration of RNA-seq (5,845 differentially expressed genes) and ChIP-seq (7,319 genes with PPARG peak) datasets resulted in the identification of 2,512 potential direct PPARG gene targets (Fig. 5D, Appendix Dataset 6). GO analyses of PPARG gene targets identified by PPARG ChIP-seq (Fig. 5E, Appendix Dataset 7) and an integrated dataset of PPARG responsive genes identified by RNA-seq of EVT cells (see Fig. 4) and PPARG gene targets identified by PPARG ChIP-seq (Fig. 5F, Appendix Dataset 8) prominently identified genes associated with positive regulation of cell projection organization and tissue migration. These represent cellular processes associated with EVT cell differentiation. In addition, GO analysis identified PPARG target genes connected to response to hypoxia, lipid transport, and WNT signaling, which are also likely important in the transition from the stem state to mature differentiated EVT cells (Fig. 5E). Evidence supporting *PLAC8*, *TFPI*, *DLX6*, *FSTL3* and *TP63*, which are known regulators of trophoblast cell differentiation (Chang *et al*, 2018; Xie *et al*, 2018; Muto *et al*, 2021; Wang *et al*, 2022; Kim *et al*, 2024), as potential direct targets of PPARG is shown (Fig. 5G). Collectively, the results support PPARG regulating genes that control development of the EVT cell lineage.

**Figure 5.**
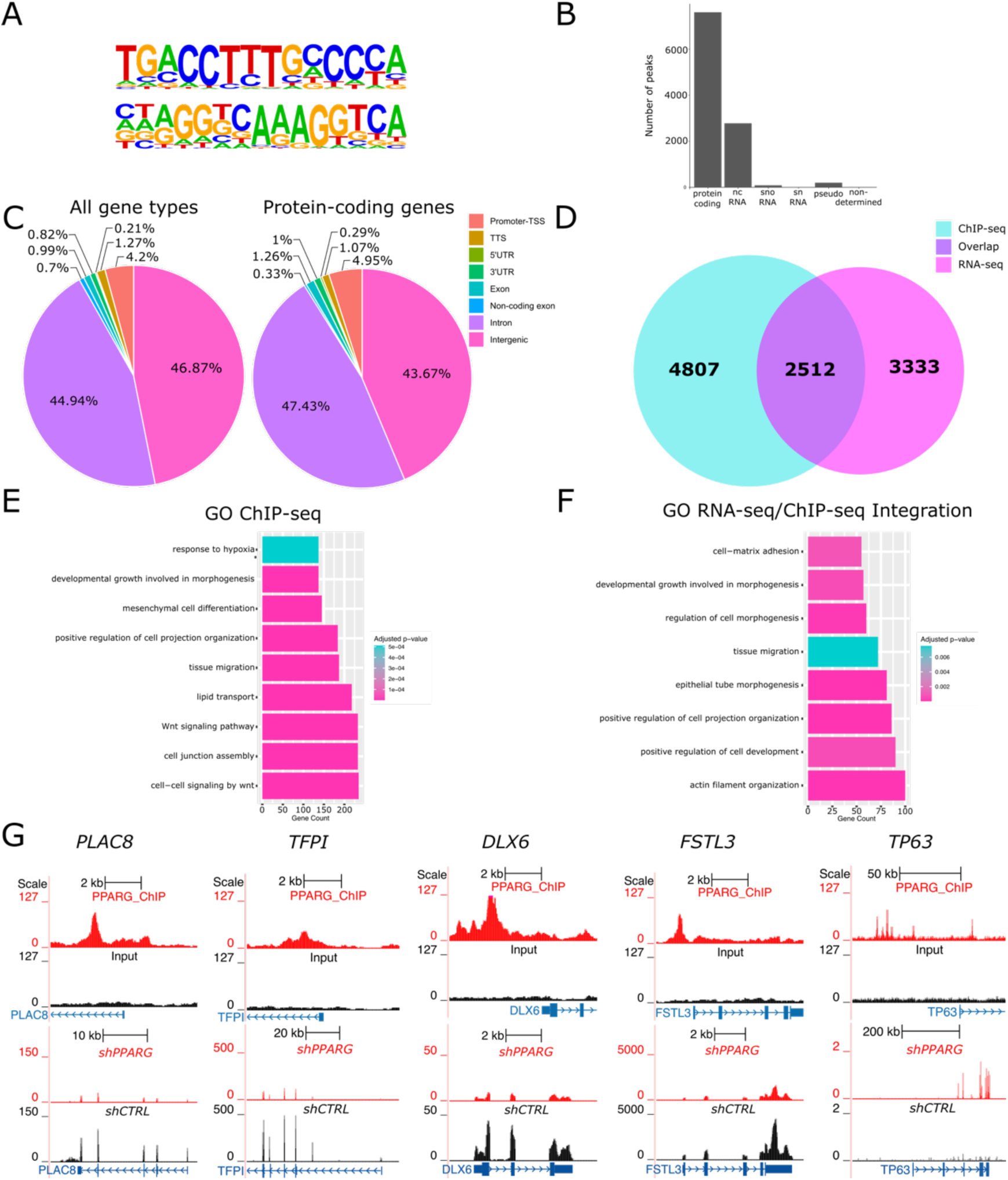
PPARG targets within the EVT cell genome identified using ChIP-seq. **(A)** Two significantly enriched PPARG binding motifs were observed within PPARG ChIP-seq peaks. **(B)** Number of peaks observed within specific locations of the genome. **(C)** Pie chart showing the distribution of PPARG peaks in genomic regions for all genes and protein-coding genes. **(D)** Venn diagram showing the integration of PPARG responsive genes identified by RNA-seq of EVT cells (**see** Fig. 4) and PPARG gene targets identified by PPARG ChIP-seq from EVT cells. **(E)** Gene ontology (**GO**) analysis of the PPARG ChIP-seq dataset performed on EVT cells. **(F)** GO analysis of an integrated dataset generated from PPARG responsive genes identified by RNA-seq of EVT cells (**see** Fig. 4) and PPARG gene targets identified by PPARG ChIP-seq from EVT cells. **(G)** Integration of PPARG ChIP-Seq peaks and RNA-seq track at selected gene loci that exhibit sensitivity to PPARG inhibition. Visualization was performed using the University of California at Santa Cruz Genome browser. Red peaks represent PPARGbound regions in the ChIP-seq and gene expression in shPPARG1 RNA-seq, whereas black peaks represent input DNA in ChIP-seq and gene expression in shCTRL RNA-seq.

### PPARG expression within the rat uterine-placental interface

The rat exhibits deep intrauterine trophoblast cell invasion (Ain *et al*, 2003), a feature of placentation also observed in the human, and has proven to be a useful in vivo model for investigating the physiological relevance of genes implicated in the regulation of the invasive trophoblast cell lineage (Chakraborty *et al*, 2016; Varberg *et al*, 2021; Muto *et al*, 2021; Kuna *et al,* 2023; Dominguez *et al,* 2024). We next sought to determine whether PPARG was expressed in trophoblast cells of the rat placenta, especially within invasive trophoblast cells of the rat uterine-placental interface. The rat placentation site is organized into three main compartments: i) labyrinth zone, ii) junctional zone, and iii) uterine-placental interface (Soares *et al*, 2012; Shukla & Soares, 2022). The labyrinth zone is comprised of trophoblast cells and fetal vasculature and represents the site of maternal-fetal exchange, whereas the junctional zone contains several trophoblast cell types, including invasive trophoblast cell progenitors, which direct pregnancy-dependent adaptations of maternal tissues. The uterine-placental interface is the uterine compartment adjacent to the placenta where invasive trophoblast cells, uterine vasculature, and maternal immune cells are situated. *In situ* hybridization was used to determine the distribution of *Pparg* transcripts. At gestation day (**gd**) 11.5, *Pparg* transcripts were localized to trophoblast cells of the developing placenta as well as to invasive trophoblast cells situated within uterine spiral arteries (Fig. 6A). The presence of *Pparg* transcripts in trophoblast cells was confirmed by co-localization with keratin 8 (*Krt8*), an epithelial cell associated transcript expressed by trophoblast cells (Fig. 6A). By gd 18.5, *Pparg* was expressed within trophoblast cells of the labyrinth zone, junctional zone, and within invasive trophoblast cells of the uterine–placental interface (Fig. 6B). Expression of *Pparg* in invasive trophoblast cells was supported by its co-localization with prolactin family 7, subfamily b, member 1 (***Prl7b1*,** Fig. 6B,C), an invasive trophoblast cell-associated transcript (Scott *et al*, 2022; Wiemers *et al*, 2003). These findings are consistent with a previously published report describing single cell RNA-seq analysis of the rat uterine–placental interface, which demonstrated that *Pparg* expression was enriched in the invasive trophoblast cell lineage (Appendix Fig. 2, Scott *et al*, 2022). *Pparg* transcript levels increased within the uterine–placental interface as pregnancy progressed (Fig. 6D). Furthermore, PPARG protein was localized to invasive trophoblast cells and throughout the junctional and labyrinth zones, as shown by immunohistochemistry (Fig. 6E). Thus, PPARG expression is a conserved feature of EVT cells of the human placenta (Fig. 1) and invasive trophoblast cells of the rat placentation site (Fig. 6).

**Figure 6.**
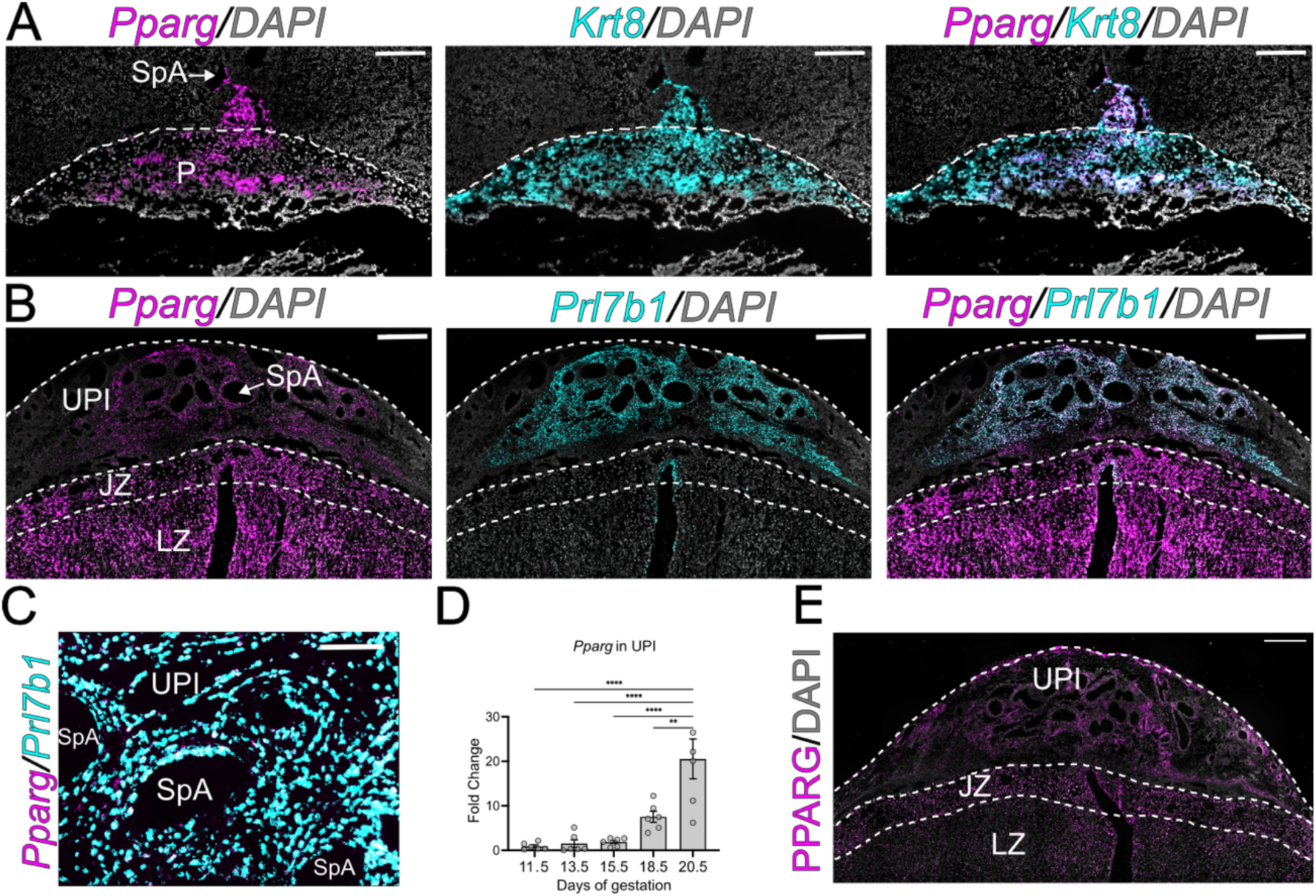
Distribution of *Pparg* transcripts and protein in rat placental tissues. *Pparg* transcripts were localized within rat placentation sites using in situ hybridization. **(A)** *Pparg* transcripts (magenta) were co-localized with *Krt8* transcripts (cyan) at gestation day (**gd**) 11.5. **(B)** Co-localization (white) of *Pparg* (magenta) and *Prl7b1* (cyan) transcripts within the gd 18.5 placentation site. **(C)** Higher magnification of the placentation site surrounding the spiral arteries showing co-localization (white) of *Pparg* (magenta) and *Prl7b1* (cyan) transcripts. **(D)** RT-qPCR measurement of *Pparg* transcript levels in uterine-placental interface tissues at gd 11.5, 13.5, 15.5, 18.5, and 20.5. Data are presented as mean ± standard error of the mean. Each data point represents a biological replicate from six independent pregnancies (n=6). Statistical analyses were performed with unpaired *t*-tests: **p<0.01, ****p<0.0001. **(E)** Immunohistochemical localization of PPARG protein within the gd 18.5 rat uterine-placental interface. Scale bars for panels A, B and E=500 μm, and panel C=50 μm. Abbreviations: SpA, spiral artery; UPI, uterine-placental interface; P, placenta; JZ, junctional zone; LZ, labyrinth zone.

### Establishment and validation of the invasive trophoblast cell-specific Pparg mutant rat

Next, we examined an in vivo role for PPARG in development of rat invasive trophoblast cells. Since, PPARG is expressed in trophoblast cell lineages throughout the rat placentation site (Fig. 6B) and global inactivation of the mouse *Pparg* gene was associated with midgestation lethality (Barak *et al*, 1999; Kubota *et al*, 1999), we utilized a conditional Cre/Lox approach to selectively disrupt *Pparg* in the invasive trophoblast cell lineage. For these experiments we used a rat model possessing Cre recombinase inserted into the *Prl7b1* locus. As noted above, *Prl7b1* is robustly and specifically expressed within invasive trophoblast cells of the rat placentation site (Wiemers *et al*, 2003; Scott *et al*, 2022; Iqbal *et al*, 2024). The *Prl7b1-Cre* specifically acts within the invasive trophoblast cell lineage without anomalous activities in non-invasive trophoblast cells or other extraembryonic and embryonic tissues (Dominguez *et al, 2024*; Iqbal *et al*, 2024). Male *Prl7b1-Cre* rats and female rats possessing *loxp* sites flanking Exons 2 and 3 of the *Pparg* gene (Fig. 7A) were used in the analysis. Specifically, *Prl7b1-Cre*/heterozygous floxed *Pparg* (***Prl7b1^Cre^/Pparg^f/+^***) males were mated with homozygous floxed *Pparg* female rats (***Pparg^f/^****^f^*) to generate *Pparg^f/f^* and *Pparg* deleted (***Pparg^d/d^***) placentation sites. Immunohistochemical analysis for PPARG demonstrated the presence of PPARG protein at the gd 18.5 uterine–placental interface of *Pparg^f/f^* but not within this compartment of *Pparg^d/d^*placentation sites (Fig. 7B).

**Figure 7.**
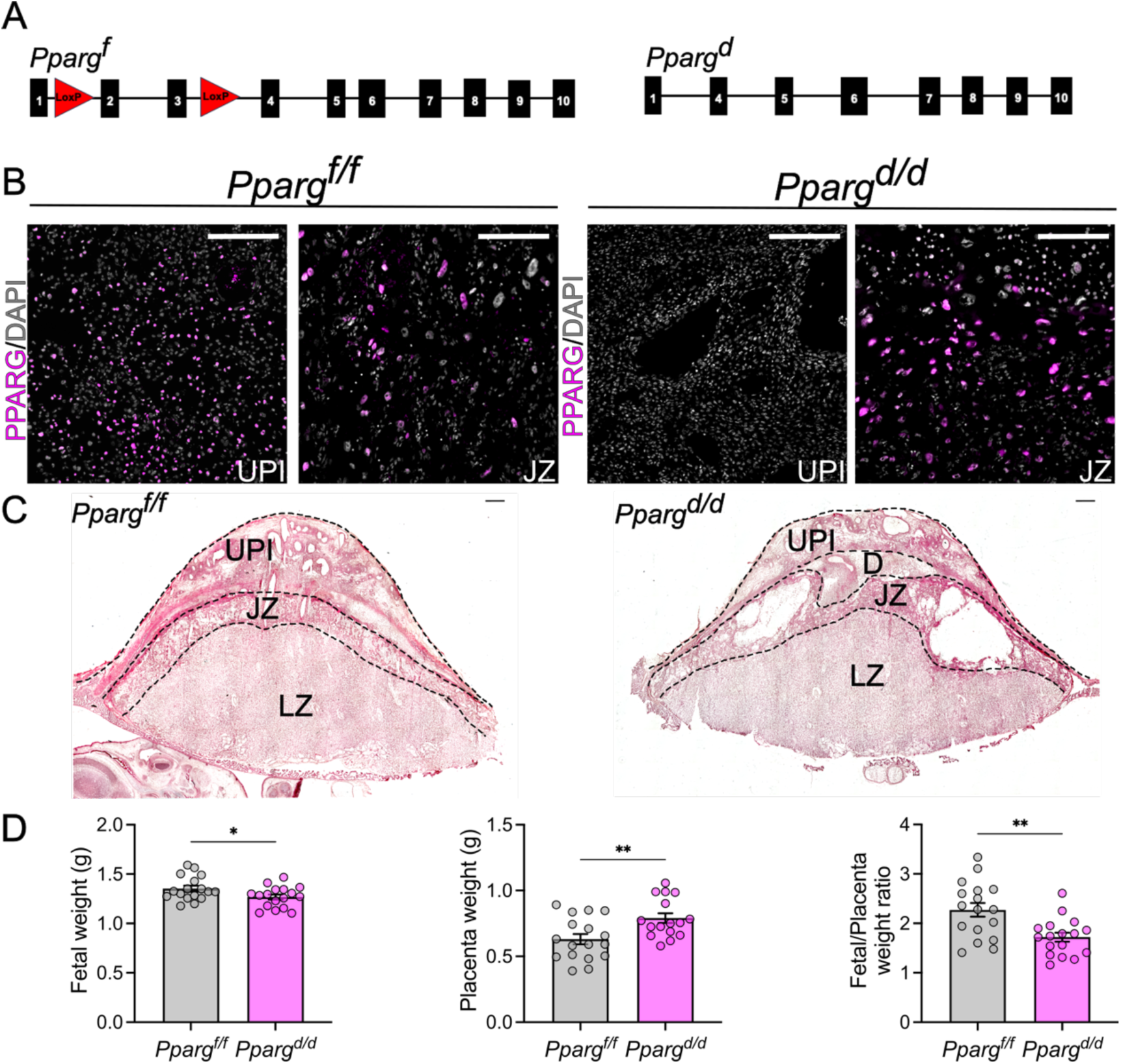
Rat invasive trophoblast-specific deletion of *Pparg*. **(A)** Schematic representation of loxP sites flanking Exons 2 and 3 of the *Pparg* gene and their conditional deletion following Cre recombinase activity driven by *Prl7b1* regulatory DNA. **(B)** PPARG immunostaining of *Pparg^f/f^* and *Pparg^d/d^* gestation day (**gd**) 18.5 placentation sites. Images show uterine-placental interface tissues (left) and junctional zone tissues (right) for each group, scale bar=50 μm. **(C)** Hematoxylin and eosin staining of *Pparg^f/f^* and *Pparg^d/d^* gd 18.5 placentation sites, scale bar=500 μm. **(D)** Placental weight, fetal weight, and fetal-to-placental weight ratio from gd 18.5 *Pparg^f/f^*and *Pparg^d/d^* conceptuses. Data are presented as mean ± standard error of the mean. Each data point represents a biological replicate from six pregnancies (*Pparg^f/f^*, n=17, *Pparg^d/d^*, n=17). Statistical analyses were performed using unpaired *t*-tests: *p<0.05, **p<0.01. Abbreviations: SpA, spiral artery; UPI, uterine-placental interface; JZ, junctional zone; LZ, labyrinth zone; D, decidua.

### PPARG involvement in rat invasive trophoblast cell lineage development and cellular dynamics within the uterine-placental interface

The structure and integrity of *Pparg^f/f^* and *Pparg^d/d^* placentation sites were different. An expanded decidual compartment was evident within *Pparg^d/d^* placentation sites (Fig. 7C). *Pparg^d/d^*placentas were larger (Fig. 7C), possessed an enlarged junctional zone (Fig. EV5) containing prominent cyst-like structures (Fig. 7C), and were less efficient than *Pparg^f/f^* placentas (Fig. 7D). Invasive trophoblast cells arise from the junctional zone (Soares *et al*, 2012; Shukla & Soares, 2022) and exhibited pronounced differences between *Pparg^f/f^* and *Pparg^d/d^* placentation sites (Fig. 8). The uterine-placental interface of *Pparg^f/f^* placentation sites was infiltrated with invasive trophoblast cells, which was not observed in *Pparg^d/d^* placentation sites, as detected by in situ hybridization for *Prl7b1* or pan-cytokeratin immunostaining (Fig. 8A,B). Deficits in intrauterine trophoblast cell invasion correlated with diminished expression of invasive trophoblast cell-associated transcripts (*Prl5a1*, *Prl7b1*, *Krt8, Krt7, Cited2, Peg3, and Tfpi*) and pan-cytokeratin immunoreactive surface area within the uterine-placental interfaces of *Pparg^d/d^* placentation sites (Fig. 8C,D; Appendix Fig. 3). Previously published single cell RNA-sequencing and single nucleus assay for transposase-accessible chromatin-sequencing datasets (Scott *et al*, 2022; Vu *et al*, 2023) were used to reveal a network of genes potentially regulated by PPARG in rat invasive trophoblast cells. These putative PPARG targets include candidate contributors to remodeling the extracellular matrix, regulation of cell movement, cytoskeletal restructuring, and transcriptional regulators that act on genes encoding other proteins affecting the invasive trophoblast cell phenotype (Appendix Table 1 and Dataset 9).

**Figure 8.**
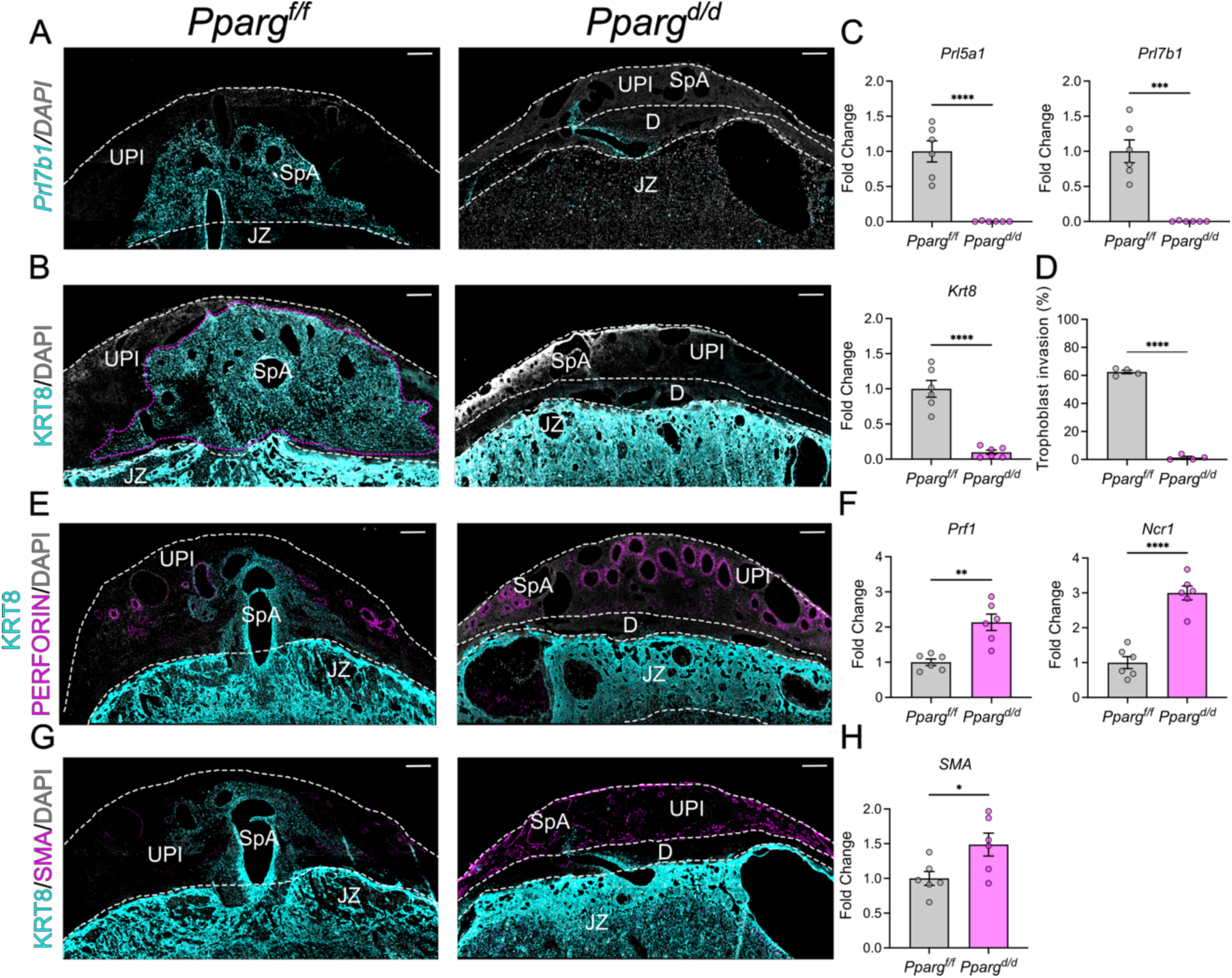
Invasive trophoblast cell-specific *Pparg* disruption impacts pregnancy-dependent transformation of the uterine-placental interface. **(A)** *In situ* hybridization showing *Prl7b1* transcripts (cyan) in gestation day (**gd**) 18.5 *Pparg^f/f^*and *Pparg^d/d^* placentation sites. **(B)** Immunohistochemical localization of KRT8 (cyan) in gd 18.5 *Pparg^f/f^* and *Pparg^d/d^*placentation sites. **(C)** RT-qPCR analysis of *Prl7b1, Prl5a1, and Krt8* expression in the uterine-placental interface of gd 18.5 *Pparg^f/f^* and *Pparg^d/d^* placentation sites (n=6). **(D)** Quantification of trophoblast cell invasion area (magenta dotted line) (n=4). **(E)** Immunohistochemical localization of perforin (**PRF1**), a natural killer (**NK**) cell-associated protein, (magenta) and KRT8 (cyan). **(F)** RT-qPCR analysis of NK cell-associated transcripts (*Prf1* and *Ncr1*) in the uterine-placental interface of gd 18.5 *Pparg^f/f^*and *Pparg^d/d^* placentation sites (n=6). **(G)** Immunohistochemical localization of smooth muscle actin (**SMA**, magenta) and KRT8 (cyan) in the uterine-placental interface of gd 18.5 *Pparg^f/f^* and *Pparg^d/d^* placentation sites. **(H)** RT-qPCR analysis of *Sma* transcript in the uterine-placental interface of gd 18.5 *Pparg^f/f^* and *Pparg^d/d^* placentation sites (n=6). Scale bars for images in panels A, B, E and G=500 μm. Data are presented as mean ± standard error of the mean. Each data point represents a distinct uterine-placental interface sample obtained from four to six pregnancies. Statistical analyses were performed using unpaired *t*-tests: \**p*<0.05, **p<0.01, \*\*\**p*<0.001, \*\*\*\**p*<0.0001. Abbreviations: UPI, uterine-placental interface; JZ, junctional zone; LZ, labyrinth zone; D, decidua; SpA, spiral artery.

Diminished intrauterine trophoblast cell invasion in *Pparg^d/d^* placentation sites was accompanied by a retention of natural killer (**NK**) cells within the uterine-placental interface (Fig. 8E) and increased expression of NK cell-associated transcripts (Fig. 8F). NK cells are known contributors to uterine spiral artery remodeling (Chakraborty *et al*, 2011; Rätsep *et al*, 2015; Renaud *et al*, 2017; Erlebacher, 2013) but were less effective than invasive trophoblast cells in disabling smooth muscle cells associated with uterine spiral arteries than invasive trophoblast cells (Fig. 8G,H).

Overall, these results highlight a key role for PPARG in directing development of the invasive trophoblast cell lineage and the important role that invasive trophoblast cells possess in regulating the cell composition and function of the uterine-placental interface.

## Discussion

Hemochorial placentation serves the important function of delivering nutrients to ensure coordinated on-time growth and development of the fetus within the female reproductive tract (Soares *et al*, 2018). This type of placentation is found in many primates and rodents, including the human and rat (Carter & Enders, 2004; Roberts *et al*, 2016). These two species exhibit extensive invasive trophoblast cell-guided uterine transformation, including restructuring of spiral arteries, which is fundamental to acquisition of maternal resources required for fetal development (Pijnenborg *et al*, 1981). We examined a role for PPARG in development of invasive trophoblast cells using human TS cells and a genetically manipulated rat model. In human placentation, PPARG was expressed in EVT cell column cells and was shown to coordinate the upregulation of genes required for cell elongation and the process of EVT cell movement. In the rat, PPARG was expressed throughout the placenta, including invasive trophoblast cells, and was essential for development of the invasive trophoblast cell lineage. Failures in PPARG-mediated gene activation also affected human TS cell proliferation and led to abnormalities within compartments of the rat placenta. The experimentation demonstrated that PPARG is a conserved regulator of invasive trophoblast cell development and function.

Earlier research connected PPARG to placentation (Schaiff *et al*, 2006; Fournier *et al*, 2011; Kadam *et al*, 2015). Insights regarding PPARG actions in placentation were unequivocally demonstrated from analysis of global PPARG null mouse models (Barak *et al*, 1999; Kubota *et al*, 1999). PPARG disruption led to midgestation lethality due to an impairment of early events in placental morphogenesis (Barak *et al*, 1999; Kubota *et al*, 1999). Considerable effort has also been directed towards analyses of roles for PPARG in regulating trophoblast cells using an assortment of in vitro models. PPARG deficient mouse TS cells exhibit impairments in proliferation and syncytiotrophoblast development and enhanced trophoblast giant cell differentiation (Parast *et al*, 2009), whereas human cytotrophoblast exposure to PPARG agonists promoted syncytiotrophoblast formation and trophoblast cell-lipid accumulation, while inhibiting trophoblast cell invasive properties (Tarrade, 2001; Tarrade *et al*, 2001; Schaiff *et al*, 2005). Our in vitro and in vivo experimentation supported an essential role for PPARG in EVT/invasive trophoblast cell lineage development. Similar observations of the involvement of PPARG in EVT cell differentiation have recently been reported (Guo *et al*, 2025). Exposure to a PPARG antagonist supported the involvement of PPARG actions in promoting EVT cell differentiation. These PPARG actions on the EVT/invasive trophoblast cell lineage differ from observations with PPARG activating ligands. PPARG can act independent of ligand and its actions can vary depending upon the ligand investigated and may differ depending on the developmental state of the target cells examined (Sauer, 2015). Thus, PPARG is a broad-spectrum regulator of placental biology with critical actions on development of the EVT/invasive trophoblast cell lineage.

As indicated above, PPARG is expressed in multiple trophoblast cell lineages (Barak *et al*, 1999; Tarrade, 2001; Fournier *et al*, 2007; Waite *et al*, 2005; Asami-Miyagishi *et al*, 2004). Each trophoblast cell lineage is responsible for controlling tasks critical for the development and function of the placenta (Soares *et al*, 2018). Context is important. PPARG actions are not only determined by its presence but also by the complement of posttranslational modifiers, heterodimerization binding partners, and co-regulators expressed by a cell (Berger & Moller, 2002; Feige *et al*, 2006; Ahmadian *et al*, 2013).

In addition to PPARG, several other proteins have been shown to contribute to the regulation of EVT cell differentiation. ASCL2, CITED2, DLX5, DLX6, EPAS1, GCM1, NOTUM, NRIP1, SNAI1, TFAP2C, TFPI, WWTR1, and ZNF439, are required for optimal transition from the TS cell stem state to EVT cells (Varberg *et al*, 2023; Shukla *et al*, 2024; Varberg *et al*, 2021; Muto *et al*, 2021; Kuna *et al,* 2023; Kim *et al*, 2024; Jeyarajah *et al,* 2022; Meinhardt *et al,* 2025). A *CRISPR* screen identified additional transcription factors contributing to the regulation of human TS cells (Shimizu *et al*, 2023). The screen demonstrated the involvement of PPARG in the control of TS cell proliferation, which was consistent with our observations, but not EVT cell differentiation. The *CRISPR* screen utilized HLA-G expression as a method for monitoring EVT cell differentiation. PPARG does not significantly affect HLA-G gene expression, which may account for differences in the outcome of the *CRISPR* screen versus our findings. Based on RNA-seq and ChIP-seq experiments we can place PPARG in a potential hierarchy of transcription factors controlling EVT cell development (Appendix Fig. 4, Varberg *et al*, 2023, 2021; Kuna *et al,* 2023; Kim *et al*, 2024; Jeyarajah *et al,* 2022; Guo *et al*, 2025). Among the regulators impacting EVT cell differentiation, ASCL2, TFPI, CITED2, TFAP2C and PPARG possess conserved actions promoting development of the invasive trophoblast cell lineage in the rat (Varberg *et al*, 2021; Muto *et al*, 2021; Kuna *et al*, 2023; Dominguez *et al*, 2024).

PPARG deficiency was associated with failed trophoblast cell invasion into the uterus leading to a dramatic reorganization of the rat placentation site. The reorganization included the retention of NK cells within the uterine-placental interface, expanded decidual tissue, and a disruption in the structural organization of the two main placental compartments (junctional and labyrinth zones). NK cells are known contributors to early events in uterine spiral artery remodeling (Chakraborty *et al*, 2011; Erlebacher, 2013; Rätsep *et al*, 2015; Renaud *et al*, 2017). NK cells vacate the uterine-placental interface as invasive trophoblast cells enter the uterus (Ain *et al*, 2003). In addition to PPARG, TFAP2C is also a driver of invasive trophoblast cell infiltration into the uterus (Dominguez *et al*, 2024). Impairments of either PPARG or TFAP2C lead to failed intrauterine trophoblast cell invasion and retention of NK cells within the uterine-placental interface. The implication is that invasive trophoblast cells are engineering the exit of NK cells from the uterine-placental interface. In the absence of invasive trophoblast cells, NK cells are supporting, at least partially, uterine spiral artery remodeling and minimizing adverse pregnancy outcomes. The nature of the signal(s) emanating from invasive trophoblast cells responsible for NK cell exodus is unknown. Cooperation of invasive trophoblast cells and NK cells at the uterine-placental interface is robust. It is expected that deficits in both maternal and extraembryonic contributions to uterine transformation would exhibit more adverse consequences. Expansion of the decidua compartment in *Pparg^d/d^* placentation sites could also be contributing to decreased trophoblast cell migration towards the uterine-placental interface within the *Pparg^d/d^* placentation sites. In addition to the uterine-placental interface, failure in the developmental progression of progenitor cells situated in the junctional zone into invasive trophoblast cells resulted in malformations within the junctional zone. These aberrations included the appearance of large cystic structures within the junctional zone and an irregular junctional zone-labyrinth zone interface. Specific cellular dysfunctions linked to PPARG deficiency underlying these structural defects in junctional zone morphogenesis are unknown.

Finally, PPARG transcriptional activity is influenced by the availability of glucose and lipids (Berger & Moller, 2002; Feige *et al*, 2006; Ahmadian *et al*, 2013). Thus, PPARG acts as a nutrient sensor. Coupling nutrient sensing with the regulation of invasive trophoblast cell transformation of the uterine vasculature would be an effective strategy for adapting to a nutrient-variable maternal environment and is testable with the in vitro and in vivo models described in this report.

## Methods and Protocols

### Human placentation site specimens

Paraffin-embedded first-trimester placental tissue specimens were obtained from the Lunenfeld-Tanenbaum Research Institute at Mount Sinai Hospital (Toronto, Canada). All tissue samples were deidentified prior to use in this study. Tissue collection was conducted following written informed consent and with approval from the Human Research Ethics Review Committees at the University of Toronto and the University of Kansas Medical Center (**KUMC**).

### Human TS cell culture

Human TS cells (CT27 and CT29) used in this study have been previously described (Okae *et al*, 2018). The TS cells originated from deidentified first trimester human placental tissue obtained from healthy women with signed informed consent and approval from the Ethics Committee of Tohoku University School of Medicine. Experimentation with human TS cells was approved by the KUMC Human Research Protection Program and the KUMC Human Stem Cell Research Oversite committee. Human TS cell lines were maintained in the stem cell state and differentiated into EVT cells as previously reported. Detailed culture conditions are provided in the Appendix section.

### Design and generation of lentiviral shRNA constructs

PPARG shRNA sequences were designed using the Genetic Perturbation Platform web portal from the Broad Institute (https://portals.broadinstitute.org/gpp/public/analysis-tools/sgrna-design), and subcloned into the pLKO.1 vector at *AgeI* and *EcoRI* restriction sites. An shRNA control (**shCTRL**, Plasmid 1864, Addgene), which does not recognize any known mammalian gene, was similarly subcloned into the pLKO.1 vector and used as a control. shRNA sequences used in these analyses are listed in Appendix section Table 2. Lentiviral packaging plasmids were obtained from Addgene and included pMDLg/pRRE (plasmid 12251), pRSV-Rev (plasmid 12253), and pMD2.G (plasmid 12259). Lentiviral particles were produced via transient transfection of the shRNA-pLKO.1 vector along with the packaging plasmids into Lenti-X cells (632180, Takara Bio USA) using Attractene (301005, Qiagen) in Opti-MEM I (51985-034, Thermo Fisher). Cells were maintained in Dulbecco’s Modified Eagle Medium (DMEM, 11995-065, Thermo Fisher) supplemented with 10% fetal bovine serum (**FBS**) for 24 h prior to supernatant collection. At that point, cells were switched to Basal Human TS Cell Medium. Conditioned medium containing viral particles was collected every 24 h over two days and stored at −80 °C until used.

### Lentiviral transduction

Human TS cells were plated at 65,000 cells per well in 6-well tissue culture-treated plates coated with 5 μg/mL collagen IV (CB40233, Thermo Fisher) and incubated for 24 h. Prior to transduction, medium was changed, and cells were incubated with 2.5 μg/mL polybrene for 30 min at 37°C. Following polybrene treatment, TS cells were transduced with 500 μL of conditioned medium containing lentiviral particles and then incubated for 24 h. Medium was changed at 24 h post transduction and selected with puromycin dihydrochloride (5 μg/mL, A11138-03, Thermo Fisher) for 2 days. Cells recovered were cultured for 1 to 3 days in Complete TS Cell Medium and then used to evaluate the effects of PPARG disruption on EVT cell differentiation.

### Cell migration assay

Trophoblast cell migration was assessed according to the method described by Shimizu and coworkers (Shimizu *et al*, 2023) with minor modifications. A suspension of 200,000 human TS cells was prepared in Matrigel (100%) and Basal Human EVT Cell Differentiation Medium (10 μL). From this mixture, 1 μL was dispensed onto a 6-well plate to form individual Matrigel drops. Six Matrigel drops were prepared. The drops were cultured in complete EVT cell differentiation medium supplemented with 0.5% Matrigel (CB-40234, Thermo Fisher) at 37°C in a humidified atmosphere containing 5% CO₂. Culture medium was replaced on day 3, and the cells were incubated for a total of five days. EVT cells migrating from the cell aggregate were identified using immunocytochemistry for HLA-G (1:300, phycoerythrin (**PE**)-labeled anti-HLA-G antibody, ab24384, Abcam). After 15 min of incubation at 37°C, cells were fixed with 4% paraformaldehyde (**PFA**). Nuclei were counterstained with 4’,6-diamidino-2-phenylindole (**DAPI**, 1:50,000, D1306, Invitrogen). Following washes with phosphate buffered saline (**PBS**, pH 7.4), the samples were imaged using a Zeiss Axio Observer 7 microscope. The area of PE-labeled signal was quantified using ImageJ software.

### Trophoblast organoids

Human TS cells were used to establish trophoblast organoids as previously described (Karvas *et al*, 2022) with minor modifications. Effects of PPARG on trophoblast organoid formation and capacity to differentiate into EVT cells were determined. Detailed culture conditions are provided in the Appendix section.

### Animals and tissue collection

Holtzman Sprague–Dawley rats were obtained from Envigo and used for in vivo experiments. Animals were housed in an environmentally controlled facility under a 14-hour light/10-hour dark lighting schedule with food and water available ad libitum. Timed pregnancies were established by mating virgin female rats (8–10 weeks of age) with adult male rats (>3 months of age). Mating was confirmed by the presence of sperm in the vaginal lavage, with the day of sperm detection considered gd 0.5. Pseudopregnant females were generated by mating with vasectomized males, and the presence of a seminal plug was designated as day 0.5 of pseudopregnancy. Pregnant rats were euthanized at different days during gestation. Some placentation sites were frozen intact in dry ice–cooled heptane and stored at –80 °C for subsequent histological analysis, while other placentation sites were dissected into uterine–placental interface, junctional zone, and labyrinth zone compartments, as previously describe (Ain *et al*, 2006).

### Conditional Pparg mutant rat

In vivo *Pparg* disruption in invasive trophoblast cells was achieved using the Cre-loxP system. LoxP cassettes were inserted into introns flanking Exons 2 and 3 of the *Pparg* locus using *CRISPR*/Cas9 mediated homologous recombination (Fig. 7, Appendix section Table 3).

*CRISPR*/Cas9 reagents and templates were electroporated into one-cell rat embryos using a NEPA21 electroporator (Nepa Gene), and embryos subsequently transferred to pseudopregnant rats. Offspring were screened for mutations via polymerase chain reaction (**PCR**), and successful loxP site insertion was confirmed by DNA sequencing (GeneWiz). A founder mutant animal was backcrossed with wild-type rats to assess germline transmission. A *Prl7b1-Cre* recombinase rat model was used to specifically disrupt *Pparg* in invasive trophoblast cells (Iqbal *et al*, 2024).

Genotyping and sex chromosome determinations were performed as previously described (Dhakal & Soares, 2017). Genomic DNA was extracted from tail-tip biopsies using the Red Extract-N-Amp Tissue PCR Kit (XNAT-1000RXN, Sigma-Aldrich) and used for genotyping. Primer sequences for detecting conditional mutations and determining sex chromosome composition are provided (Appendix section Table 4).

### RT-qPCR

Total RNA was isolated from tissues using TRIzol reagent (15596018, Thermo Fisher). One μg of RNA was used to synthesize complementary DNA (**cDNA**) using the High-Capacity Reverse Transcription Kit (4368814, Applied Biosystems), followed by a 1:10 dilution in nuclease-free water. Quantitative PCR (**qPCR**) was performed using PowerSYBR Green PCR Master Mix (4367659, Thermo Fisher) and transcript-specific primer sets (250 nM). Primer sequences are provided in Appendix section, Tables 5 and 6. qPCR was conducted with a QuantStudio 5 Real-Time PCR System (Thermo Fisher) using the following cycling conditions: an initial denaturation step at 95 °C for 10 min, followed by 40 cycles of two-step PCR (95°C for 15 sec, 60°C for 1 min). A dissociation curve was then generated with the following steps: 95°C for 15 sec, 60°C for 1 min, and a final at 95°C for 15 sec. Relative mRNA expression levels were calculated using the ΔΔCt method. Glyceraldehyde 3-phosphate dehydrogenase (*Gapdh*) and RNA polymerase II subunit A (*POLR2A*) were used as reference genes for rat and human samples, respectively.

### In situ hybridization

In situ hybridization was performed using paraffin-embedded human placenta and frozen rat placentation site tissue sections and the RNAscope® Multiplex Fluorescent Reagent Kit v2 (Advanced Cell Diagnostics) following the manufacturer’s instructions. Custom-designed probes were used to detect the following transcripts: human *PPARG* (441691, NM_138712.3, target region: 699–1779), human *NOTUM* (430311, NM_178493.5, target region: 259–814), rat *Pparg* (313251, NM_001145367.1, target region: 305–1932); rat *Krt8* (87304-C2, NM_199370.1, target region: 134–1472), and rat *Prl7b1* (860181-C2, NM_153738.1, target region: 28–900). Nuclei were visualized using DAPI (323108, Advanced Cell Diagnostics). Images were captured using a Zeiss Axio Observer 7 microscope.

### Immunohistochemistry of placental tissues

Formalin fixed paraffin-embedded human placental tissue sections and frozen rat placentation site sections (10 μm thick) fixed in 4% paraformaldehyde were processed for immunohistochemical analyses. Tissue sections were blocked with 10% goat serum (50062Z, Thermo Fisher) and incubated overnight at 4°C with primary antibodies against vimentin (1:300, sc-6260, Santa Cruz Biotechnology), pan-cytokeratin (1:100, F3418, Sigma-Aldrich), perforin (1:300, TP251, Amsbio), or PPARG (human: 1:100, sc-7273, Santa Cruz Biotechnology; rat: 1:100, 2435S, Cell Signaling Technology). Immunostaining was visualized using fluorescence-tagged secondary antibodies: Alexa Fluor 488-conjugated goat anti-mouse IgG (A11001, Thermo Fisher) and Alexa Fluor 568-conjugated goat anti-rabbit IgG (A11011, Thermo Fisher). Sections were counterstained with DAPI, mounted in Fluoromount-G (0100-01, Southern Biotech), and imaged using a Nikon 90i upright microscope equipped with Photometrics CoolSNAP-ES monochrome camera (Roper). Regions of the uterine–placental interface containing cytokeratin-positive cells were quantified using ImageJ software, as previously described (Dominguez *et al*, 2024; Rosario *et al*, 2008).

### Immunohistochemistry of trophoblast organoids

Organoids were recovered from Matrigel using Cell Recovery Solution (354253, Corning) and dissociated by gentle pipetting. After 30 min incubation on ice, samples were centrifuged (600 xg, 5 min), washed with PBS, and fixed in 4% paraformaldehyde on ice for 30 min. Fixed organoids were suspended in 20% sucrose solution and incubated at 4°C overnight. The sucrose solution was subsequently replaced with 1 mL of warm gelatin/sucrose solution and incubated at 37°C for 15 min. Blocks of gelatin containing trophoblast organoids were froze for 1-2 min and stored at-80°C until processed for cryosection. Sections were permeabilized in PBS containing 4% bovine serum albumin and 0.5% Triton X-100. Primary antibodies to HLA-G (1:300, phycoerythrin (**PE**)-labeled, ab24384, Abcam), CGB (1:500, ab9528, Abcam) and TP63 (1:500, ab124762, Abcam) were incubated with tissue sections overnight at 4°C, followed by secondary antibody incubation. Sections were counterstained with DAPI, mounted in Fluoromount-G (0100-01, Southern Biotech), and imaged using a Nikon 90i upright microscope equipped with Photometrics CoolSNAP-ES monochrome camera (Roper).

### Western blotting

Human trophoblast cell lysates were processed using the radioimmunoprecipitation assay lysis buffer (sc-24948A, Santa Cruz Biotechnology) and protein concentrations determined using the DC Protein Assay Kit (5000112, Bio-Rad). Proteins (50 to 70 µg/lane) were separated by sodium dodecyl sulfate polyacrylamide gel electrophoresis. Separated proteins were electrophoretically transferred to polyvinylidene difluoride membranes (10600023, GE Healthcare) for 1 h at 25 V on a semi-dry transfer apparatus (Bio-Rad). Membranes were subsequently blocked with 5% milk for 1 h at room temperature, followed by incubation with antibodies to PPARG (1:200, 2435S, Cell Signaling Technology or GAPDH (1:5,000, AM4300, Invitrogen) in 5% milk overnight at 4°C. After primary antibody incubation, the membranes were washed in Tris-buffered saline with Tween 20 (**TBS-T**) three times (10 min/wash) at room temperature. The membranes were then incubated with anti-mouse immunoglobulin G conjugated to horseradish peroxidase (**HRP**; 1:5000, 7076, Cell Signaling Technology) or anti-rabbit immunoglobulin G conjugated to horseradish peroxidase (**HRP**; 1:5000, 5127, Cell Signaling Technology) in 5% milk for 1 h at room temperature, washed in TBS-T three times (10 min/wash) at room temperature, immersed in Immobilon Crescendo Western HRP Substrate (WBLUR0500, Sigma-Aldrich), and luminescence detected using Chemi Doc MP Imager (Bio-Rad). The integrated densities of proteins separated by western blotting were quantified using NIH ImageJ. PPARG protein intensities were normalized to GAPDH protein intensities. All quantifications were performed on at least three independent biological replicates.

### RNA-seq analysis

Transcript profiles were generated from control and PPARG shRNA treated human CT27 TS cells following EVT cell differentiation. Three independent lentiviral transductions for each shRNA were analyzed. RNA was isolated and integrity assessed using an Agilent 2100 Bioanalyzer. cDNA libraries prepared with Illumina TruSeq RNA preparation kits according to the manufacturer’s instructions. Barcoded cDNA libraries were multiplexed onto a TruSeq paired-end flow cell and sequenced (100-bp paired-end reads) with a TruSeq 200-cycle sequencing-by-synthesis kit. Samples were sequenced on an Illumina NovaSeq platform at the KUMC Genome Sequencing Facility. Reads from *.fastq files were mapped to the human reference genome (GRCh37) using CLC Genomics Workbench 12.0 (Qiagen). Transcript abundance was expressed as total reads, and a false discovery rate of 0.05 was used as a cutoff for significant differential expression. Statistical significance was calculated by empirical analysis of digital gene expression followed by Bonferroni’s correction.

### ChIP-seq analysis

PPARG ChIP was performed on human CT27 TS cells following EVT cell differentiation using commercially available SimpleChIP^®^ Enzymatic Chromatin Immunoprecipitation Kit (Magnetic Beads, 9003, Cell Signaling Technology) and PPARG antibody (2435S, Cell Signaling Technology). Three independent replicates were processed. The ChIP-seq library was constructed using the Tecan Ovation Ultralow DNA System V2 (NuGEN Technologies, Redwood City, CA) and sequenced on an Illumina NovaSeq platform. Details of data processing and analyses are presented in Supplemental Material.

### Statistics

Statistical analyses were performed with GraphPad Prism 10.2.3 software. Statistical comparisons were evaluated using Student’s *t* test or one-way analysis of variance with Dunnett’s post hoc test. Statistical significance was determined as P<0.05.

### Study approval

All protocols using rats were approved by the University of Kansas Medical Center Animal Care and Use Committee, Kansas City, Kansas (approved protocol 22-01-220).

## Supporting information

Expanded View

## Data availability

RNA-seq and ChIP-seq datasets are available at the Gene Expression Omnibus database, https://www.ncbi.nlm.nih.gov/geo/ (GSE302291 and GSE302293).

The source data of this paper are collected in the following database record, https://www.ebi.ac.uk/biostudies/bioimages/studies/S-BIAD2948

All data generated and analyzed in this study are included in the published article and supporting files.

## Author Contributions

E.M.D and M.J.S. conceived and designed the research; E.M.D., A.M-I. and A.F. performed experiments; K.I., H.O., and T.A. provided experimental tools for analysis; E.M.D., A.M-I., K.C., G.T., M.P., and M.J.S. analyzed the data and interpreted results of experiments; E.M.D. and M.J.S. prepared the manuscript. All authors read, contributed to editing, and approved the final version of manuscript.

## Acknowledgments

We thank Brandi Miller and Stacy Oxley for administrative assistance. The work was supported by the Lalor Foundation (E.M.D., A.M.-I.), Kansas Idea Network of Biomedical Research Excellence, P20 GM103418 (E.M.D, A.M.-I.), Program of the Joint Usage/Research Center for Developmental Medicine and High Depth Omics, IMEG, Kumamoto University (H.O.), NIH grants: HD115834 (A.M.-I.), HD104071 (K.I.), HD020676 (M.J.S.), HD105734 (M.J.S.), HD112559 (M.J.S., G.T.), the Sosland Foundation (M.J.S.), and the Donald C. Johnson Research Endowment Fund.

