## Supplementary material for "PPARG directs trophoblast cell fate and establishment of the uterine-placental interface": Expanded View

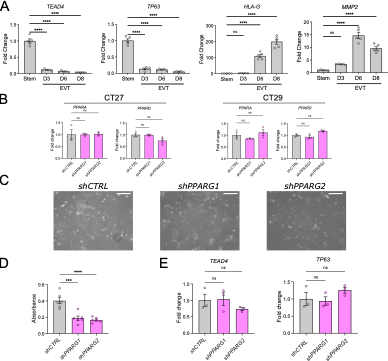

**Figure Expanded View 1. Involvement of PPARG on the behavior of CT27 (X,X) trophoblast stem (TS) cells.** **(A)** Expression of TEAD4 and TP63 (stem-state-associated transcripts), and HLA-G and MMP2 (EVT cell-associated transcripts) in CT27 TS cells in the stem state (day 0, D0) and on day 3 (D3), day 6 (D6) and day 8 (D8) of EVT cell differentiation (n=5). **(B)** PPARA and PPARD transcript levels in CT27 and CT29 TS cells after 8 days of EVT differentiation in control versus PPARG1 and PPARG2 shRNA-treated conditions (n=3). **(C)** Phase-contrast images of TS cells in the stem state expressing control or PPARG1 and PPARG2 shRNAs (scale bar=100 µm). **(D)** Quantification of cell proliferation by crystal violet staining in control versus PPARG1 and PPARG2 shRNA-treated stem-state TS cells (n=6). **(E)***TEAD4* and *TP63* transcript levels in control versus PPARG1 and PPARG2 shRNA-treated TS cells in the stem state (n=3). RT-qPCR was used to measure transcript levels. Data are presented as mean ± standard error of the mean. Each data point represents a biological replicate. Statistical analyses were performed using one-way analysis of variance followed by Dunnett’s post hoc test: ***p<0.001, ****p<0.0001.

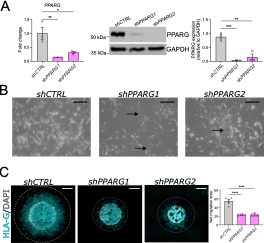

**Figure Expanded View 2. Involvement of PPARG in the differentiation of CT29 (X,Y) trophoblast stem (TS) cells to extravillous trophoblast (EVT) cells**. **(A)** PPARG transcript and protein levels in CT29 TS cells after 8 days of EVT differentiation in control versus PPARG1 and PPARG2 shRNA-treated conditions (n=3). **(B)** Phase-contrast images of EVT-differentiated cells expressing control or PPARG1 and PPARG2 shRNAs (scale bar=100 µm). Arrows indicate cells exhibiting stem-like morphology. **(C)** HLA-G immunofluorescence detection of migrating cells (scale bar=500 µm) and corresponding quantification of net migration area during EVT cell differentiation in control versus PPARG1 and PPARG2 shRNA-treated cells (n=6). RT-qPCR was used to measure transcript levels. Data are presented as mean ± standard error of the mean. Each data point represents a biological replicate. Statistical analysis was performed using one-way analysis of variance followed by Dunnett’s post hoc test : *p<0.05, **p<0.01, ***p<0.001, ****p<0.0001.

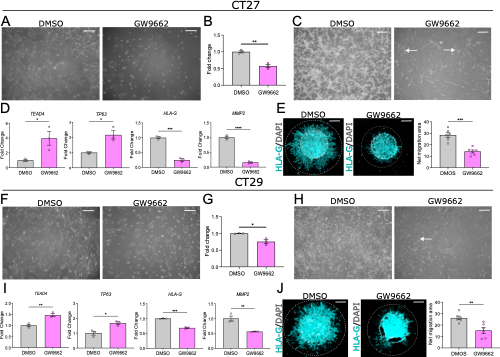

**Figure Expanded View 3. Effects of a PPARG antagonist (GW9662) on EVT cell differentiation**. **(A)** Phase-contrast images of CT27 TS cells maintained in the stem state and treated with DMSO (vehicle) or GW9662 (10 µM), scale bar=100 µm. **(B)** Quantification of cell numbers in DMSO and GW9662 (10 µM) treated CT27 TS cells. **(C)** Phase-contrast images of CT27 TS cells exposed to EVT cell differentiation conditions and treated with DMSO and GW9662 (10 µM), scale bar=100 µm. **(D)** RT-qPCR measurements of TEAD4 and TP63 (stem-state-associated transcripts) and HLA-G and MMP2 (EVT cell-associated transcripts) in DMSO or GW9662 (10 µM) treated CT27 TS cells following 8 days of EVT cell differentiation. **(E)** HLA-G immunofluorescence detection of migrating CT27 TS cells (scale bar=500 µm) and corresponding quantification of net migration area during EVT cell differentiation in DMSO versus GW9662 (10 µM) treated cells (n=6). **(F)** Phase-contrast images of CT29 TS cells maintained in the stem state and treated with DMSO (vehicle) or GW9662 (10 µM), scale bar=100 µm. **(G)** Quantification of cell numbers in DMSO and GW9662 (10 µM) treated CT29 TS cells. **(H)** Phase-contrast images of CT29 TS cells exposed to EVT cell differentiation conditions and treated with DMSO or GW9662 (10 µM), scale bar=100 µm. **(I)** RT-qPCR measurements of TEAD4 and TP63 (stem-state-associated transcripts) and HLA-G and MMP2 (EVT cell-associated transcripts) in DMSO or GW9662 (10 µM) treated CT29 TS cells following 8 days of EVT differentiation. **(J)** HLA-G immunofluorescence detection of migrating CT29 TS cells (scale bar=500 µm) and corresponding quantification of net migration area during EVT cell differentiation in DMSO or GW9662 (10 µM) treated cells (n=6). Data are presented as mean ± standard error of the mean. Each data point represents a biological replicate. Statistical analysis was performed using unpaired *t*-tests: *p<0.05, **p<0.01, ***p<0.001, ****p<0.0001.

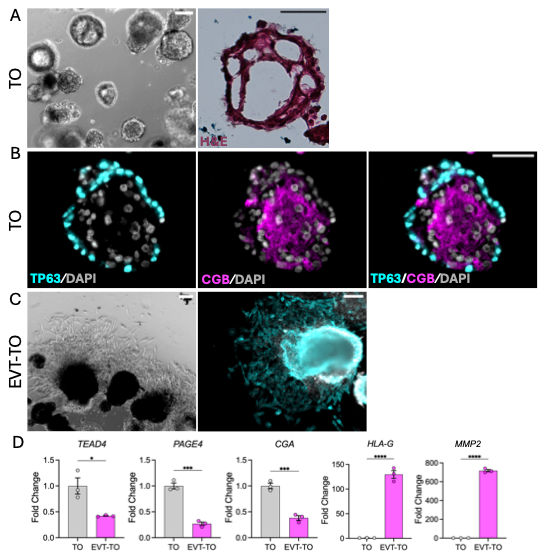

**Figure Expanded View 4. Characterization of TS cell derived trophoblast organoids (TO).** **(A)** Phase contrast image and hematoxylin and eosin of TO, scale bar=100 and 50 μm. **(B)** TP63 (cyan) and CGB (magenta) immunodetection in TO, scale bar=50 μm. **(C)** HLA-G immunocytochemistry in TO transferred to EVT cell differentiation conditions (**EVT-TO**), scale bar=50 and 100 μm. **(D)** RT-qPCR of stem, syncytiotrophoblast and EVT associated transcripts in TO and EVT-TO (n=3). Data are presented as mean ± standard error of the mean. Each data point represents a biological replicate. Statistical analysis was performed using unpaired *t*-tests: *p<0.05, ***p<0.001, ****p<0.0001.

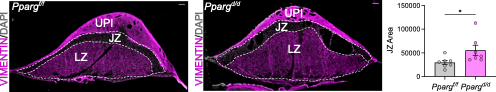

**Figure Expanded View 5. Junctional zone areas in *Pparg^f/f^* and *Pparg^d/d^* placentation sites.** **Left and central panels**, Immunohistochemical staining for vimentin, which is positive in the labyrinth zone (**LZ**) and uterine-placental interface (**UPI**) and negative in the junctional zone (**JZ**). **Right panel**, quantification of JZ areas in *Pparg^f/f^* and *Pparg^d/d^* placentation sites. Data are presented as mean ± standard error of the mean. Each data point represents a biological replicate from six independent pregnancies (n=7). Statistical analyses were performed with an unpaired *t*-test: *p<0.05.
